# Distinct contributions of cohesin-STAG1 and cohesin-STAG2 to the pluripotent and totipotent states

**DOI:** 10.64898/2026.11.13.766432

**Authors:** Maria Solé-Ferran, Daniel Giménez-Llorente, Miriam Rodríguez-Corsino, Ana Cuadrado, Sergi Aranda, Luciano di Croce, Ana Losada

**Affiliations:** Chromosome Dynamics group, Molecular Oncology Programme, Spanish National Cancer Research Centre (CNIO), Melchor Fernández Almagro 3, 28029 Madrid, Spain; Centre for Genomic Regulation (CRG), Barcelona Institute of Science and Technology, Dr. Aiguader 88, 08003 Barcelona, Spain

**Keywords:** cohesin, pluripotency, totipotency, differentiation, 3D genome

## Abstract

Cohesin organizes the three-dimensional genome and thereby regulates gene expression. In somatic cells, cohesin complexes carry either STAG1 or STAG2, paralogs that impart distinct chromatin binding dynamics and are both essential for mouse embryonic development. How each variant contributes to the maintenance and transitions of pluripotent cell state remains poorly understood. Using degrons and knockout clones in mouse embryonic stem cells, we show that neither STAG1 nor STAG2 is individually required for differentiation towards primitive endoderm or for transitions between ground-state and metastable pluripotency. Nonetheless, STAG2 is necessary to support naïve pluripotency in the metastable but not in the ground-state. Depletion of either STAG protein upregulates totipotency-associated transcripts, yet an increased proportion of two-cell-like cells is only observed upon STAG1 loss. These findings suggest complementary roles for the two cohesin variants in stabilizing the pluripotent state. While cohesin-STAG1 prevents return to totipotency, cohesin-STAG2 restrains exit towards commitment.

## Introduction

Cohesin is a highly conserved protein complex that is essential for sister chromatid cohesion (Losada et al. 1998). It is also a major organizer of the three-dimensional genome thanks to its ability to extrude DNA loops (Nishiyama 2019; Davidson and Peters 2021). In somatic cells, cohesin consists of SMC1A, SMC3, RAD21 and the STAG subunit, which can be STAG1 or STAG2 (Cuadrado and Losada 2020). Additional regulators modulate its ability to form and extrude loops (NIPBL) and to be retained or released from chromatin (PDS5, WAPL, SORORIN), the latter being also dependent on SMC3 acetylation by ESCO1/2 (reviewed by Alonso-Gil and Losada 2023a; Solé-Ferran and Losada 2025). The two STAG proteins share extensive sequence homology, and either one is sufficient for cellular viability. However, both are required for normal embryogenesis (van der Lelij et al. 2017; Remeseiro et al. 2012; De Koninck et al. 2020). Complexes carrying STAG1 or STAG2 (hereafter, cohesin-STAG1 or cohesin-STAG2) display distinct chromatin-binding properties and some differences in genomic distribution (Kojic et al. 2018; Wutz et al. 2020; Cuadrado et al. 2022; Alonso-Gil et al. 2023). Cohesin-STAG1 is found at CTCF-bound sites and contributes to demarcation of Topological Associated Domains (TADs). It shows increased residence time on chromatin, preferential association with CTCF and acetylated SMC3 and reduced dependence on NIPBL. Cohesin-STAG2, in contrast, can also be found at CTCF-independent genomic elements, associates preferentially with WAPL, and is more dynamic and dependent on NIPBL. The molecular determinants of these distinct behaviors remain to be clarified, although an AT-hook motif present in the N-terminus of STAG1 appears to be responsible for some of these features (Sakata et al. 2025).

Despite the central role of cohesin in shaping genome architecture, its acute depletion results in modest effects on steady state transcription, with only a subset of enhancer-promoter (E-P) interactions depending on cohesin-mediated loop extrusion (Rao et al. 2017; Thiecke et al. 2020; Kim et al. 2026; Hansen et al. 2026; Aboreden et al. 2026; Popay et al. 2026). It has been proposed that cohesin could be of particular importance when cells must change their transcriptional programs to differentiate or respond to stimuli (Cuartero et al. 2018; Minderjahn et al. 2022; Robles-Rebollo et al. 2022). Indeed, changes in chromatin conformation accompany transcriptional changes during stem cell differentiation and development, and several reports provide evidence for the contribution of cohesin (Bonev et al. 2017; Liu et al. 2023; Lupianez et al. 2015; Kiefer et al. 2023; Mahara et al. 2024). In accordance, germ line mutations in cohesin or its regulators are at the origin of human developmental syndromes, known as cohesinopathies (Kline et al. 2018; Solé-Ferran and Losada 2025).

Pluripotent mouse embryonic stem cells (mESC) represent an attractive system to investigate the role of cohesin on regulation of steady-state and inducible gene expression programs. Maintenance of naïve pluripotency requires a precise control of transcriptional networks by master regulators such as OCT4, SOX2 and NANOG, whose expression is supported by large superenhancer domains enriched for Mediator, p300 and cohesin (Hnisz et al. 2013; Whyte et al. 2013). When cultured in serum-LIF medium, mESC adopt a metastable state characterized by heterogeneous expression of naïve pluripotency factors and display some features that resemble the primed post-implantation epiblast, such as higher DNA methylation and lineage priming (Chambers et al. 2007; Bernstein et al. 2006; Hackett and Azim Surani 2014). In contrast, culture in the presence of MEK and GSK3 inhibitors (also known as “2i”) promotes naïve ground-state pluripotency, more closely resembling the pre-implantation blastocyst, with a more homogenous expression of pluripotency genes, reduced DNA methylation, and diminished lineage priming (Hackett and Azim Surani 2014; Ying et al. 2008). Perturbation of cohesin or its regulators has been shown to disrupt superenhancer organization and reduce the expression of pluripotency-associated genes (Apostolou et al. 2013; Rao et al. 2017; Liu et al. 2021).

In addition to pluripotent cells, mESC cultures contain a rare population of cells (∼0.1- 1%) that transiently acquire totipotent features. Totipotency is restricted to the early embryo and is linked to zygotic genome activation (ZGA). In the mouse embryo, ZGA occurs in two waves: a minor wave beginning at the late zygote stage is followed by a more extensive major wave at the late 2-cell stage (Abe et al. 2015; Aoki 2022). The rare cells mentioned above recapitulate some of the molecular features of totipotent 2-cell embryo and are therefore termed 2-cell-like cells (2CLC). These features include activation of repetitive elements, such as MERVL, expression of totipotency-associated genes (e.g., *Zscan4*), global DNA hypomethylation, and widespread chromatin relaxation (Macfarlan et al. 2012; Genet and Torres-Padilla 2020). Knockdown of cohesin subunits (*Smc1 or Rad21*), WAPL or CTCF results in upregulation of totipotency-associated genes and increases the percentage of this population of 2CLC (Macfarlan et al. 2012; Zhu et al. 2021; Kriz et al. 2021; Olbrich et al. 2021; Pezic et al. 2023; Carey et al. 2026).

Previous results have shown that the relative abundance of cohesin-STAG1 and cohesin-STAG2 changes when mESC transition from ground-state to metastable pluripotency, with STAG2 levels increasing and STAG1 levels declining (Cuadrado et al. 2019; Pezic et al. 2023). Quantitative immunofluorescence of preimplantation embryos also shows a different regulation of the maternal contribution of the two proteins as well as their differential abundance in the two lineages emerging at blastocyst stage, the trophectoderm and the pluripotent inner cell mass (Solé-Ferran et al. 2026). Downregulation of either variant in mESC has different consequences for genome architecture and gene expression, with cohesin-STAG2 being located and regulating the chromatin contacts at superenhancer regions and promoting expression of pluripotency genes (Cuadrado et al. 2019). Whether these differences are maintained in ground-state pluripotency, and how mESC transition between the two naïve pluripotency states or exit pluripotency to acquire a new identity in the presence of a single cohesin variant remains unknown.

Here, we have addressed those questions by analyzing the behavior of mESC expressing degron-tagged versions of STAG proteins or *knocked out* (KO) for the corresponding genes. We find that loss of STAG1 or STAG2 neither prevents mESC from exiting pluripotency to acquire a new cellular identity nor affects the transitions between ground-state and metastable pluripotency. Cohesin-STAG2 is important for proper expression of pluripotency genes in the metastable state, but not in the ground-state. We further report that the repression of totipotency-associated genes in mESC is sensitive to perturbation of cohesin dynamics including loss of any of the two cohesin variants, but emergence of 2CLC is only observed in the absence of cohesin-STAG1. Together, these results reveal distinct and context-dependent contributions of cohesin-STAG1 and cohesin-STAG2 to transcriptional states associated with totipotency and pluripotency.

## Results

### Loss of STAG1 or STAG2 reduces chromatin-bound cohesin levels but does not affect cell proliferation

To assess the effects of STAG1 or STAG2 loss on stem cell differentiation, mESC clones in which these proteins could be acutely depleted were generated. For that, a FKBP12^F36V^ degron tag (hereafter abbreviated FKBP) was fused to the N-terminus of STAG1 or the C-terminus of STAG2. Reduced levels of the corresponding STAG protein compared to the parental cells were sometimes observed even in the absence of dTAG^V^-1 addition (hereafter, dTAG), suggesting that the tag may affect protein stability. In any case, cellular levels of STAG1 or STAG2 dropped below 10% one hour after exposure to dTAG (Figure 1A-B; Supplementary Figure 1). To assess whether the effects of depleting one or the other variant are due to a specific contribution of STAG1 or STAG2, or to a global decrease in cohesin, a heterozygous clone carrying the FKBP degron tag on one of the *Rad21* alleles was generated, which allowed partial reduction of RAD21 levels (Figure 1B). As a complementary experimental system, *Stag1* KO and *Stag2* KO clones were also obtained, in which the respective proteins were undetectable by immunoblotting (Figure 1C).

**Figure 1.**
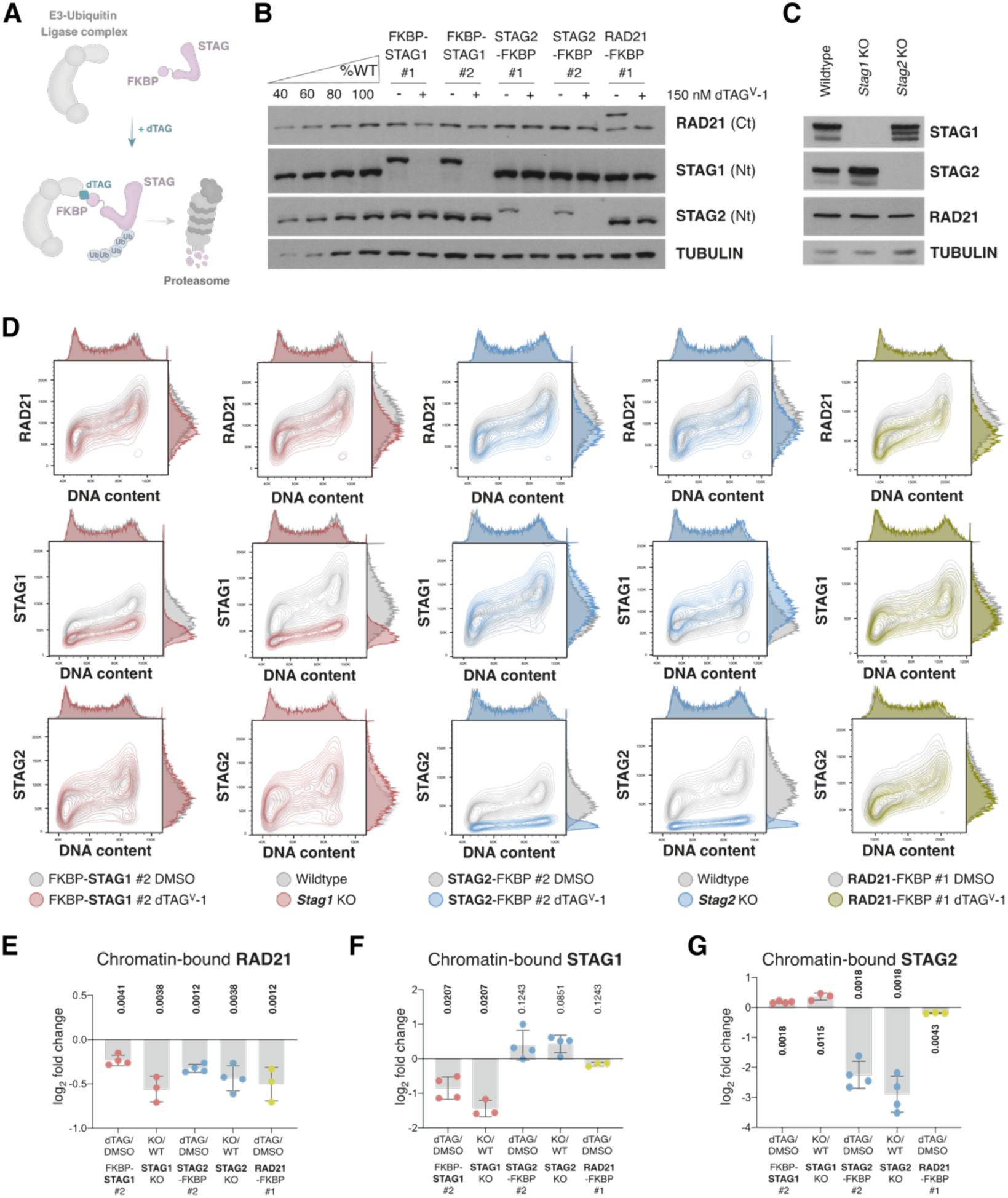
Similarly reduced cohesin levels in STAG1 and STAG2 depleted cells. A. Illustration of the dTAG system. B. Immunoblot of whole cell extracts from parental wildtype (WT) cells, FKBP-STAG1, STAG2-FKBP and RAD21-FKBP clones at 24 hours of treatment with 150nM dTAG^v^-1 (+) or DMSO (-). TUBULIN was used as a loading control. A ladder with increasing amounts of WT cell extract was used for semiquantitative assessment of remaining protein levels. C. Immunoblot of whole cell extracts from WT, *Stag1* KO and *Stag2* KO clones. TUBULIN was used as a loading control. D. Contour plots of chromatin-bound levels of the indicated cohesin subunits along the cell cycle assessed by flow cytometry. For each clone, dTAG^v^-1-treated or KO clones (colored) are compared to DMSO-treated or WT cells (in grey), respectively. E-H. Quantification of at least three independent experiments as in (D) for RAD21 (E), STAG1 (F) and STAG2 (G). The log2 fold change relative to controls was calculated from the median fluorescence intensity for each independent experiment (dot), the bar represents the mean and the whiskers the standard deviation. To assess statistical significance, multiple paired-tests were performed on the raw median fluorescence intensity values, with correction for multiple comparisons using the two-stage Bejamini, Krieger and Yekutieli false discovery rate procedure. P-values below 0.05 are highlighted in bold. See also Supplementary Figure 2.

To further characterize all these clones, we next measured chromatin-bound levels of cohesin using a quantitative flow cytometry assay (Alonso-Gil and Losada 2023b). Efficient depletion (dTAG) or absence (KO) of the targeted STAG variant was accompanied by a compensatory increase of the remaining paralog, but this increase was insufficient to maintain the same amount of cohesin on chromatin as in the non-depleted cells, as evidenced by the decrease in RAD21 (Figure 1D-H). Importantly, the reduction of RAD21 was similar in all conditions including the RAD21 heterozygous clone, validating its inclusion in the subsequent experiments. We also examined the chromatin levels of cohesin regulators PDS5A and WAPL, which showed a trend towards reduction in both *Stag1* and *Stag2* KO lines as well as in cells depleted of RAD21 for 24 hours, but not in STAG1 or STAG2 depleted cells (Supplementary Figure 2). NIPBL, CTCF, and acetylated SMC3 were unchanged in all conditions (Supplementary Figure 2). These results suggest that the cohesin complex is in excess relative to these cofactors and therefore, changes in the ratio of the complex and its cofactors in the different conditions tested are unlikely to underlie the consequences that loss of STAG1 or STAG2 may have in the behavior of mESC.

Given the relevance of cohesin for many nuclear functions, we also checked cell cycle progression and cell proliferation. Cell cycle distribution was unaffected by the loss of STAG1 or STAG2 (Supplementary Figure 3A-B). Analysis of cellular proliferation in *Stag1* and *Stag2* KO clones by cell counting revealed no significant differences (Supplementary Figure 3C). Proliferation in degron clones was assessed by flow cytometry, monitoring the progressive dilution of a fluorescent dye over three consecutive days. No significant differences in proliferation were observed between control and dTAG-treated cells across all clones tested (Supplementary Figure 3D-F). Reduction of RAD21 levels in the RAD21-FKBP clone exposed to dTAG resulted in a modest decrease in the percentage of cells in S-phase, although proliferation remained unaffected (Supplementary Figure 3B and 3E-D). These results are important to rule out that potential effects in differentiation experiments are an indirect consequence of altered proliferation.

### Either STAG variant is sufficient to acquire primitive endoderm identity

To address the importance of cohesin-STAG1 and cohesin-STAG2 in pluripotency exit and acquisition of a more committed cell identity, we differentiated mESC to primitive endoderm (PrE) cells which, after thirteen days, have a high expression of PDGFRa and form colonies with a characteristic cobblestone morphology (Supplementary Figure 4A-C). Such colonies were observed after differentiation of the degron cell lines both in the presence and absence of dTAG, and in WT and KO clones (Figure 2A). Consistently, western blot analyses confirmed that cells in all conditions efficiently downregulated the pluripotency marker NANOG and upregulated the PrE-marker GATA6 (Figure 2B). The efficiency of PrE differentiation was quantified by measuring the proportion of PDGFRa-positive cells in each culture by flow cytometry (Figure 2C-D). Although this proportion varied among individual clonal lines, no substantial differences were observed between DMSO- and dTAG-treated degron clones or between *Stag1* KO or *Stag2* KO and wildtype mESC. Upregulation of the PrE transcripts *Gata6*, *Sox17* and *Foxa2* was also detected in all tested conditions, confirming acquisition of the PrE fate (Supplementary Figure 4D-F). Thus, either cohesin-STAG1 or cohesin-STAG2 is sufficient for mESC to exit pluripotency and acquire the new transcriptional and morphological identity of PrE cells.

**Figure 2.**
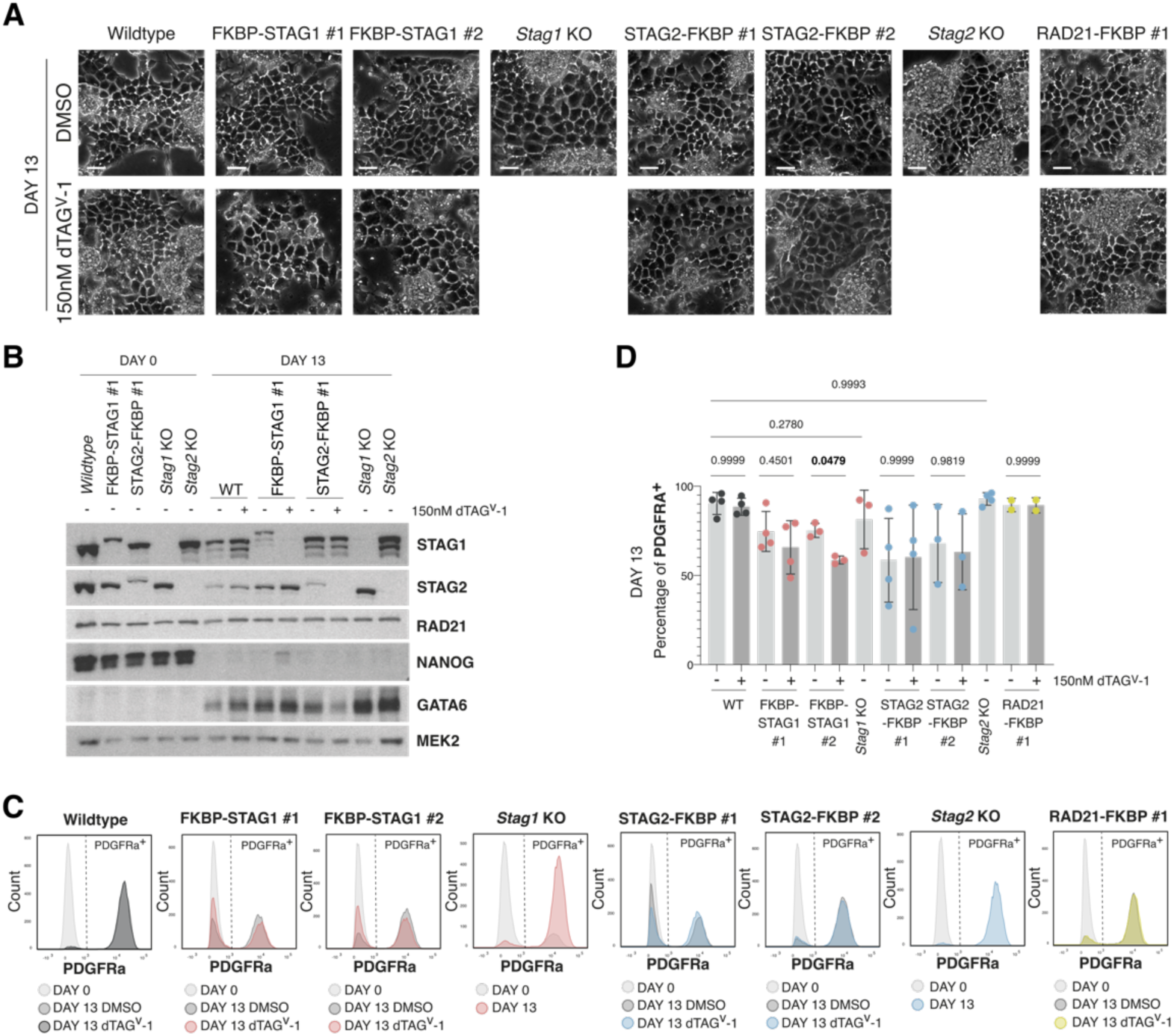
STAG1-or STAG2-deficient mESC can successfully differentiate to Primitive Endoderm. A. Representative images of DMSO or dTAG treated mESC and clones differentiated towards PrE. Scale bar, 40µm. B. Whole cell extracts from DAY 0 and DAY 13 of the differentiation protocol were analyzed with the indicated antibodies. MEK2 was used as a loading control. C. Representative histogram plots of PDGFRA intensity measured by flow cytometry D. Percentage of PDGFRA positive cells measured by flow cytometry as in C. Each dot corresponds to one experiment; the bars represent the mean and the whiskers the standard deviation. P-values< 0.05 are highlighted in bold. They were calculated using a two-way ANOVA followed by Sídák correction for multiple comparisons. Each comparison was between dTAG- and DMSO-treated cells or between KO and WT cells.

### Transitions between pluripotent states occur in the absence of STAG1 or STAG2

Next, we asked whether mESC can properly transition between metastable and ground-state naïve pluripotency in the absence of either STAG variant. Cells grown in serum-LIF were first switched to 2i for four days. The tightly packed but irregular colonies typical of serum-LIF cultures turned to compact, dome-shaped colonies with well-defined borders in both DMSO- and dTAG-treated degron clones (Figure 3A-B; Supplementary Figure 5A). Quantitative RT-PCR analyses further showed that complete ablation of STAG1 or STAG2, or partial depletion of RAD21, did not impair transcriptional changes driving this transition, including upregulation of pluripotency gene *Nanog* and downregulation of primed pluripotency genes *Dnmt3b* and *Otx2* (Figure 3C-E). Next, we examined the transition from a ground state to a metastable state by withdrawing 2i from media for four days (Figure 3F), during which the reverse changes are expected. All degron clones grown in the presence of DMSO or dTAG were able to complete the transition, as judged by visual inspection of colony morphologies, and transcriptional changes in *Nanog, Dnmt3b* and *Otx2* (Figure 3G-J; Supplementary Figure 5B). These results indicate that mESC lacking cohesin-STAG1 or cohesin-STAG2 retain the ability to transition between the two pluripotent states. However, although the transcriptional changes occurred in the expected direction, STAG2-deficient cells consistently exhibited a mild decrease in pluripotency gene expression together with a stronger induction of lineage-priming genes in comparison to their corresponding controls (Figure 3H-J). Moreover, if cells were passaged and cultured for two additional days in serum-LIF, flattened colonies typical of differentiating cells could be observed in at least one of the two FKBP-STAG2 clones in the presence of dTAG, in contrast to its control (Figure 3G). These findings suggest that cohesin-STAG2 contributes to maintaining the balance between pluripotency and lineage priming.

**Figure 3.**
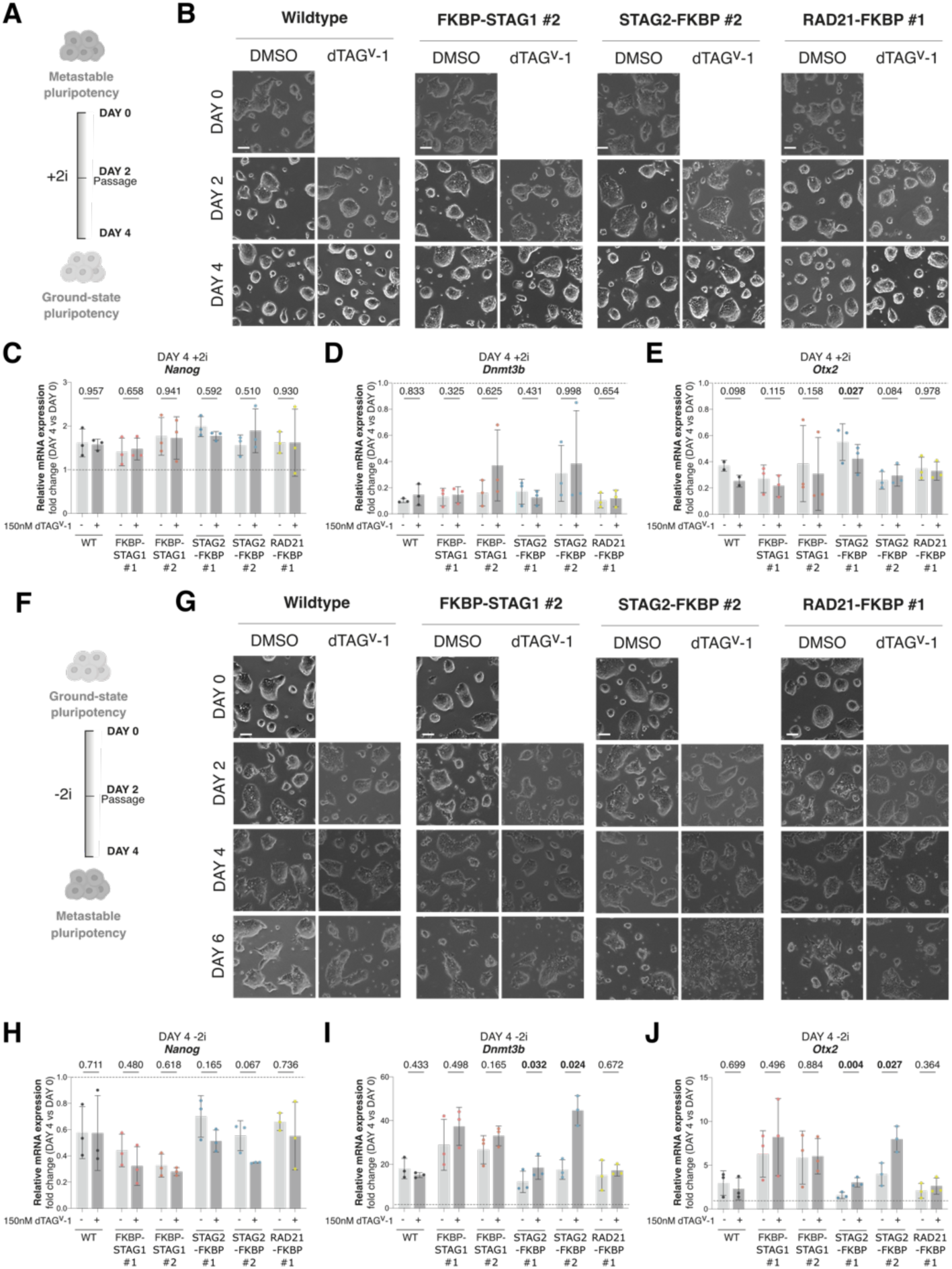
Successful transitions between ground-state and metastable pluripotency in the absence of STAG1 or STAG2. A. Experimental design for the transition between metastable and ground-state pluripotency. B. Representative images of cells of the indicated clones on days 0, 2 and 4 of the transition as in (A). Scale bar, 20µm. Images of the remaining clones can be found in Supplementary Figure 5A. C-E. Fold changes in *Nanog* (C), *Dnmt3b* (D) and *Otx2* (E) transcripts in DMSO or dTAG^v^-1 treated clones at transition DAY 4 from (B). Each sample was first normalized by *Gapdh* expression and then to expression at DAY 0 (dashed line for reference). Each dot represents the fold change from one experiment; the bar represents the mean and the whiskers the standard deviation. P-values < 0.05 are highlighted in bold. They were calculated using a two-way ANOVA followed by Tukey multiple-comparisons test on the raw ΔCt values. Each comparison was between dTAG-treated cells and DMSO-treated controls. F-J. As in A-E for the transition between ground-state and metastable pluripotency.

### Maintenance of metastable, but not ground-state pluripotency requires cohesin-STAG2

Consistent with the results for STAG2-FKBP clone #2 six days after 2i withdrawal (Figure 3G), 72-hour dTAG treatment of degron clones routinely maintained in serum-LIF consistently induced morphological changes only upon STAG2 degradation (Figure 4A). A modest but consistent reduction in NANOG protein levels (Figure 4B-C) as well as downregulation of *Nanog* and *Klf4* transcripts (Figure 4D-E) was also observed after ablation of STAG2. In contrast, STAG1 degradation did not significantly reduce pluripotency gene expression (Figure 4B-E). Interestingly, degradation of either STAG variant in ground-state (2i) condition, in which pluripotency genes are more homogeneously and highly expressed, did not affect their expression at either the transcript or protein level, nor did it induce detectable morphological changes (Figure 4C-F; Figure 5A). We therefore conclude that cohesin-STAG2, but not cohesin-STAG1, is required for maintenance of the metastable pluripotent state. Despite this, both *Stag1* KO and *Stag2* KO mESC clones can be grown in both serum-LIF and 2i conditions (Supplementary Figure 6A) and express pluripotency genes at levels comparable to those of wildtype cells (Supplementary Figure 6B). It is likely that adaptive mechanisms upon long-term growing without STAG2 allow the *Stag2* KO clone to preserve functional pluripotency.

**Figure 4.**
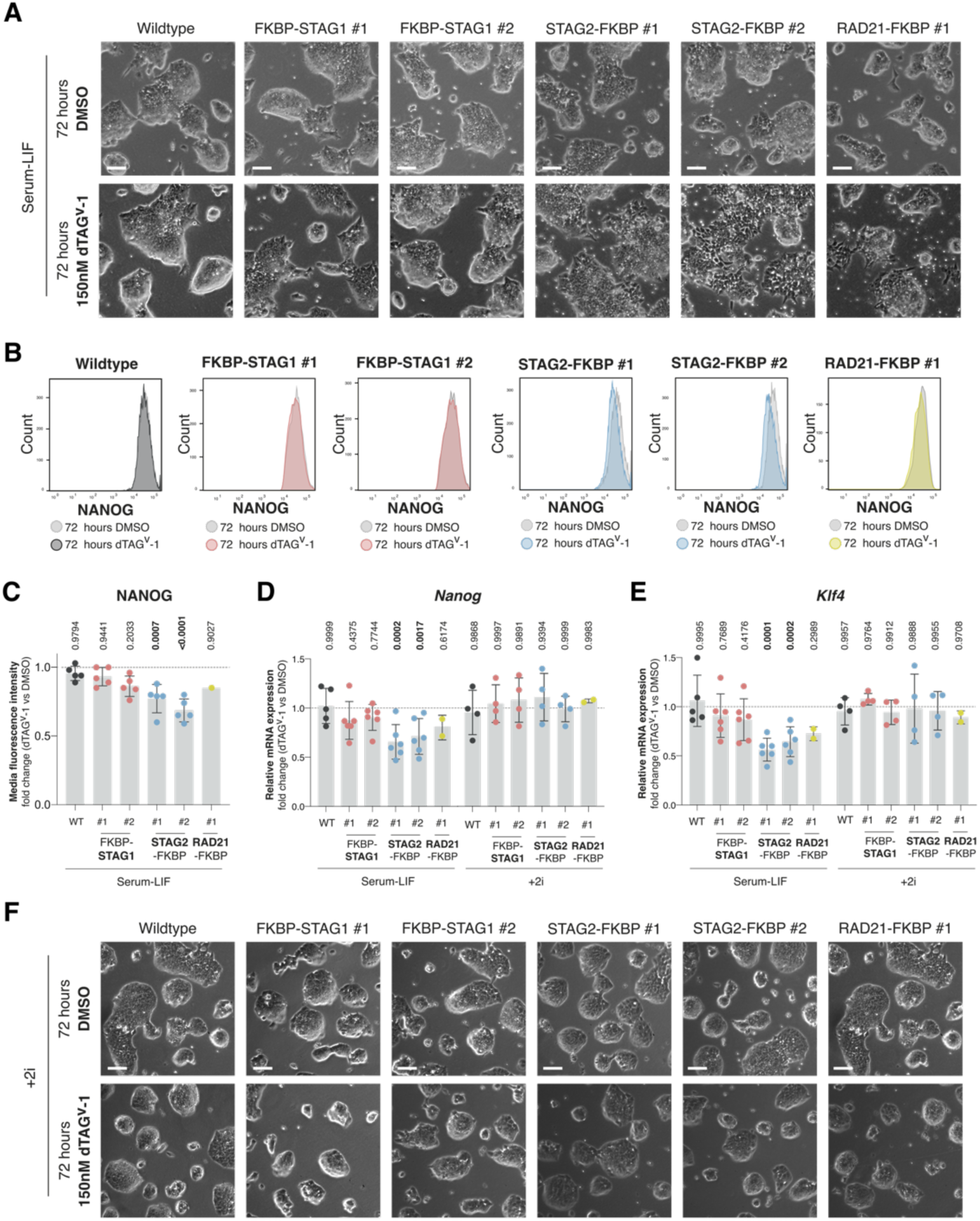
Cohesin-STAG2 is required to maintain metastable but not ground-state pluripotency. A. Representative images of wildtype (WT) mESC or the indicated clones treated with DMSO or dTAG^v^-1 for 72 hours in serum/LIF (metastable) medium. Scale bar, 20µm. B. Representative histogram plots of NANOG intensity measured by flow cytometry in cells treated with DMSO or dTAG^v^-1 for 72 hours. C. Quantification of NANOG median fluorescence intensity from (B). For each clone, the fold change was calculated relative to its DMSO-treated control (dashed line for reference). Each dot represents the fold change calculated from one independent experiment, the bar represents the mean and the whiskers the standard deviation. P-values were calculated on the raw median fluorescence intensity values using a two-way ANOVA followed by Sídák correction for multiple comparisons. D-E. Fold changes in *Nanog* (D) or *Klf4* (E) transcripts after 72 hours of DMSO or dTAG^v^-1 treatment in cells grown in serum/LIF (from A) or 2i (from F), as indicated. Each dot represents the fold change relative to its DMSO-treated control (dashed line for reference) from one experiment; the bar represents the mean and the whiskers the standard deviation. P-values were calculated using a two-way ANOVA followed by Sídák correction for multiple comparisons on the raw ΔCt values. Each comparison was between dTAG-treated cells and DMSO-treated controls. P-values< 0.05 are highlighted in bold in C-E. F. Representative images of wildtype (WT) mESC or the indicated clones treated with DMSO or dTAG^v^-1 for 72 hours in 2i-containing medium (ground-state). Scale bar, 20µm.

**Figure 5.**
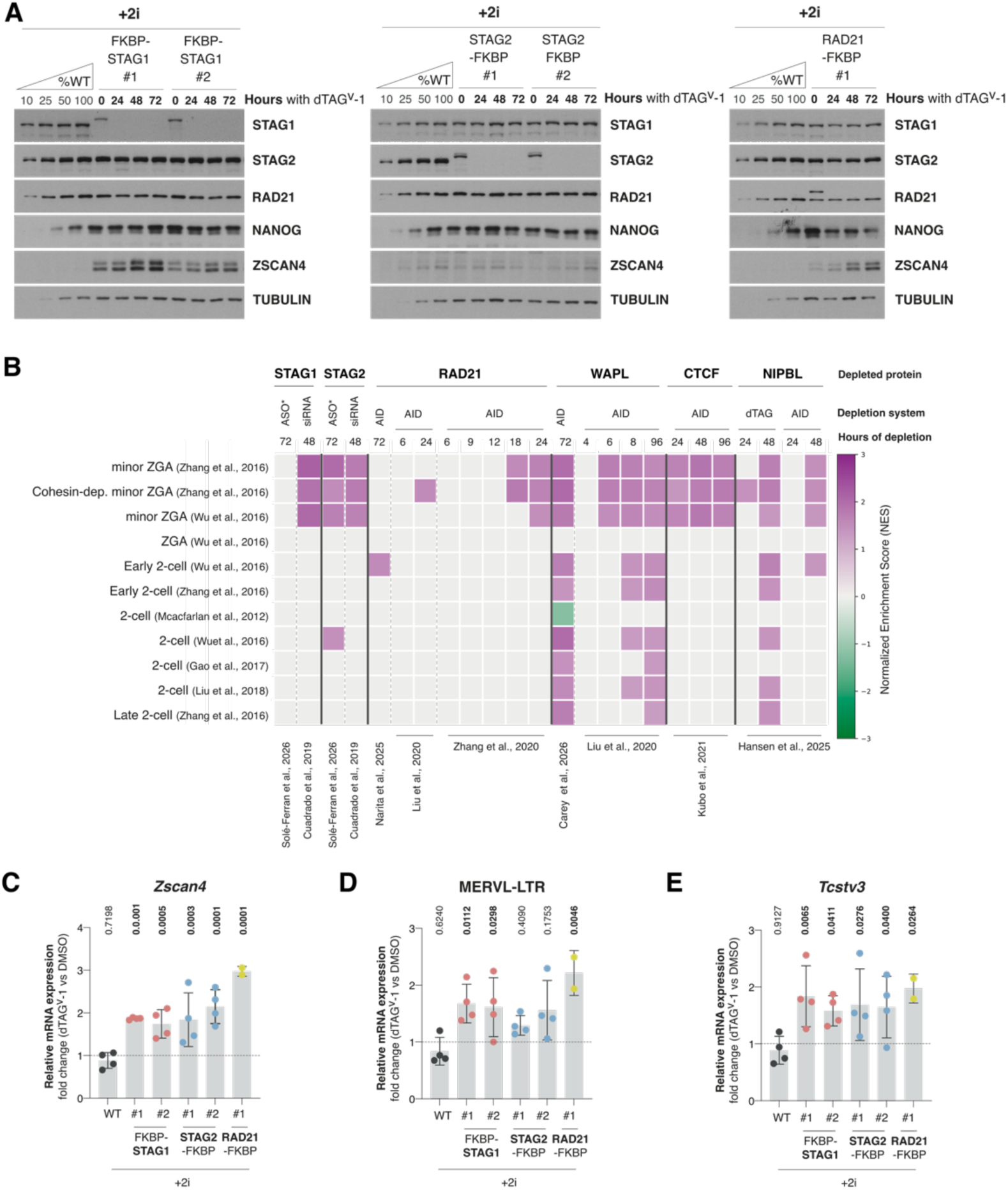
Perturbation of cohesin or its regulators results in upregulation of totipotency-related transcripts. A. Representative immunoblots of whole cell extracts from the indicated clones at different times of dTAG addition in cells cultured with 2i. TUBULIN was used as a loading control. A ladder with increasing amounts of WT cell extract allows for quantitative comparisons. B. Heatmap showing normalized enrichment scores (NES) from gene set enrichment analysis (GSEA) across multiple gene expression datasets from cells in which cohesin subunits or regulators were depleted as indicated on top. Each row represents an individual gene set. Colored cells indicate significant NES enrichment (FDR < 0.05) while non-significant comparisons are shown in grey. Antisense oligonucleotides (ASO*) were injected in mouse embryos at zygote stage and transcription was analyzed three days later at blastocyst stage (columns 1 and 3). All other perturbations were performed in mESC. “Cohesin-dep. minor ZGA genes” (second row from top) are a subset of minor ZGA genes that were described to be cohesin-dependent (Zhang et al., 2020). C-E. Fold changes in *Zscan4* (C), MERVL-LTR (D) or *Klf4* (E) transcripts after 72 hours of DMSO or dTAG treatment in cells grown in 2i. Each dot represents the fold change relative to its DMSO-treated control (dashed line for reference) from one experiment, the bar represents the mean and the whiskers the standard deviation. P-values< 0.05 are highlighted in bold. They were calculated using a two-way ANOVA followed by Sídák correction for multiple comparisons on the raw ΔCt values. Each comparison was between dTAG-and DMSO-treated cells.

### Cohesin dynamics regulate totipotency-associated gene expression

It has been suggested that cohesin depletion facilitates the transition from pluripotency to totipotency in mESC (Zhu et al., 2021). Consistent with this possibility, protein levels of ZSCAN4, a hallmark of 2CLC, raised in a time-dependent manner following dTAG addition to STAG1-FKBP and RAD21-FKBP clones growing in 2i (Figure 5A). To further explore this issue, we reanalyzed published gene expression datasets from mESC or early embryos in which cohesin subunits or regulators had been depleted (Figure 5B). We identified upregulation of minor-ZGA genes following siRNA-mediated depletion of either STAG1 or STAG2 in mESC (Cuadrado et al. 2019). In mouse blastocysts obtained from zygotes microinjected with antisense oligonucleotides to deplete either STAG protein, upregulation of minor-ZGA genes was observed following STAG2 loss (Figure 5B; (Solé-Ferran et al. 2026)). Acute RAD21 depletion in mESC also induced expression of minor-ZGA genes, and it occurred in a time-dependent manner, consistent with our observations following immunoblot analysis (Figure 5A-B; (Zhang et al. 2020; Narita et al. 2025; Liu et al. 2021). The same was true for CTCF and WAPL, in line with previous findings (Figure 5B; (Liu et al. 2021; Kubo et al. 2021; Carey et al. 2026)). Analysis of WAPL depletion reveals a time-dependent upregulation of these genes, with a robust induction of 2-cell-associated genes becoming evident 8 hours after eliciting degradation. Similarly, increased expression of minor-ZGA and 2-cell genes was evident 48 hours after triggering NIPBL elimination but not at earlier times (Figure 5B;(Hansen et al. 2026)). These results support a contribution of cohesin to the regulation of the totipotent state and further suggest that this contribution is not direct, as the consequences of its perturbation are not immediate.

### Cohesin-STAG1 represses the emergence of 2-cell-like cells

As indicated above, ZSCAN4 protein levels clearly rose after degradation of STAG1 or RAD21, while the increase was less evident for STAG2 degradation (Figure 5A). This could be in part because basal levels of ZSCAN4, in the absence of dTAG, were already higher in the FKBP-STAG1 and RAD21-FKBP clones compared to wildtype or STAG2-FKBP cells (Figure 5A; compare wildtype to 0 hours). The reason for this difference is unclear. Analysis of transcript levels revealed upregulation of *Zscan4* and two other totipotency genes (*MERVL-LTR* and *Tcstv3*) in all five clones 72 hours after dTAG addition (Figure 5C). To determine whether these transcriptional changes reflect an increased proportion of 2CLC in the culture, we performed high-throughput confocal imaging of ZSCAN4 and MERVL-gag. As positive control, we treated wildtype mESC with NaCl, as osmotic stress has been shown to promote the transition to the totipotent state (Canat et al., 2023). Consistent with the immunoblot results (Figure 5A), we observed a higher basal frequency of ZSCAN4+ and MERVL-gag+ cells in FKBP-STAG1 and RAD21-FKBP clones compared to wildtype and STAG2-FKBP cells (Figure 6). Importantly, this proportion increased after (total) STAG1 and (partial) RAD21 depletion but remained unchanged after STAG2 degradation. Thus, cohesin-STAG1 plays a more relevant role than cohesin-STAG2 in the repression of the totipotent state.

**Figure 6.**
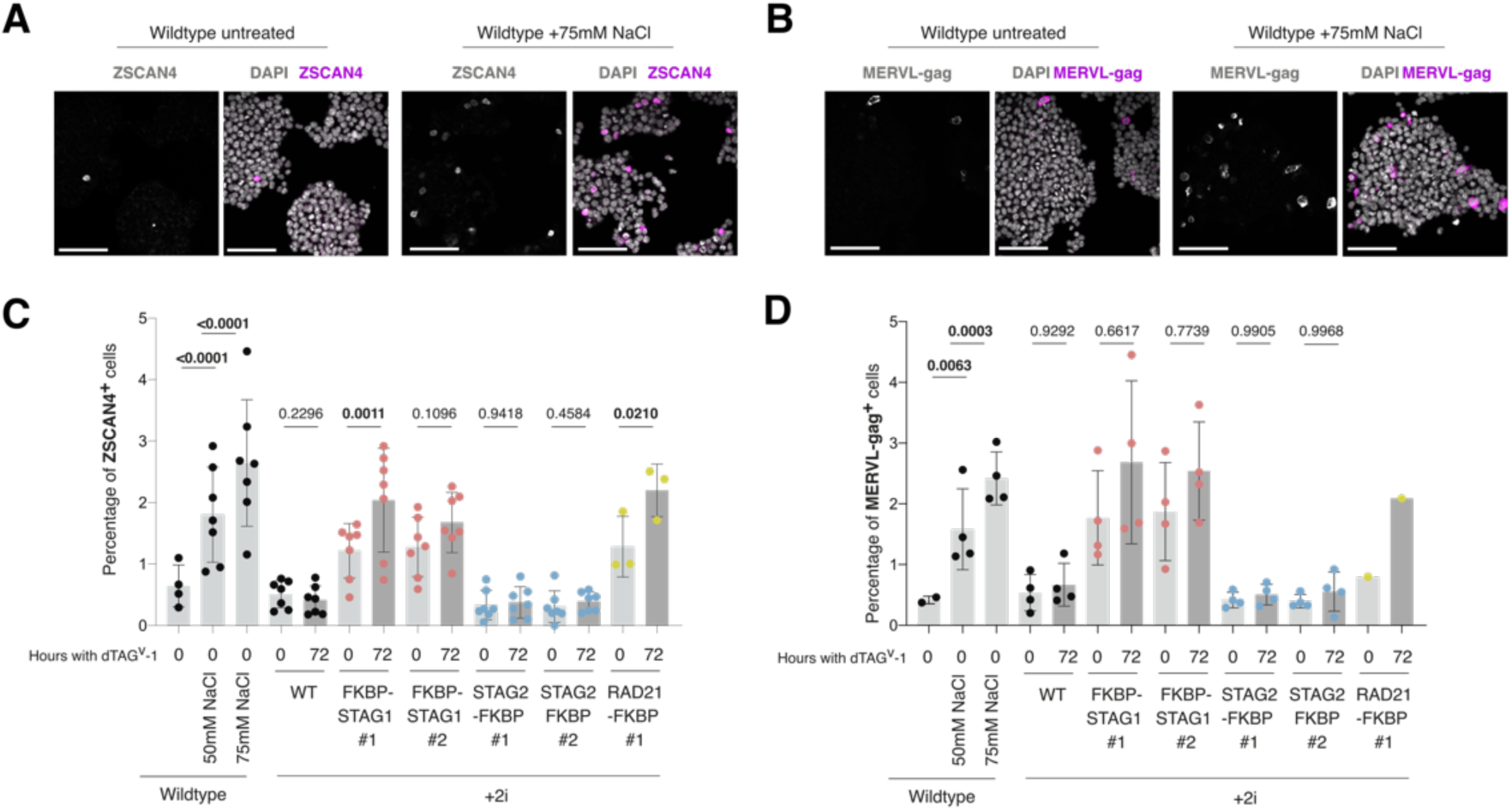
Emergence of 2CLC among 2i-grown mESC upon cohesin reduction. A-B. Representative immunofluorescence images of wildtype mESC untreated or treated with 75mM NaCl for 18 hours followed by 6 hours of recovery, stained with ZSCAN4 (A) or MERVL-gag (B). DAPI used as nuclei staining. Scale bar, 100µm. C-D. Quantification of the percentage of ZSCAN4 and MERVL-gag positive cells from (A) and (B), respectively, in the indicated conditions. Each dot represents the percentage of positive cells in one experiment, the bar represents the mean and the whiskers the standard deviation. To assess statistical significance, the proportions of positive cells were logit-transformed, then a mixed-effects model was applied followed by Sídák multiple-comparisons test. P-values< 0.05 are highlighted in bold.

## Discussion

Here, we have explored the contributions of cohesin-STAG1 and cohesin-STAG2 to maintenance and exit of pluripotency. We found that either one is sufficient to support exit from pluripotency and acquisition of a new cellular identity, as well as the transitions between the two naïve pluripotency states. In contrast, cohesin-STAG2 is specifically required to maintain the proper balance between pluripotency and commitment in metastable mESC, while being dispensable for ground-state pluripotency. Also, cohesin-STAG1 is more relevant for restricting 2CLC emergence in ground-state cell cultures. Thus, both cohesin variants contribute to maintaining the pluripotent state through distinct yet complementary actions. Cohesin-STAG1 prevents return to totipotency and cohesin-STAG2 restrains exit towards commitment (Figure 7). Taken together, our results identify cohesin dynamics as an important determinant of the balance between pluripotency and totipotency.

**Figure 7.**
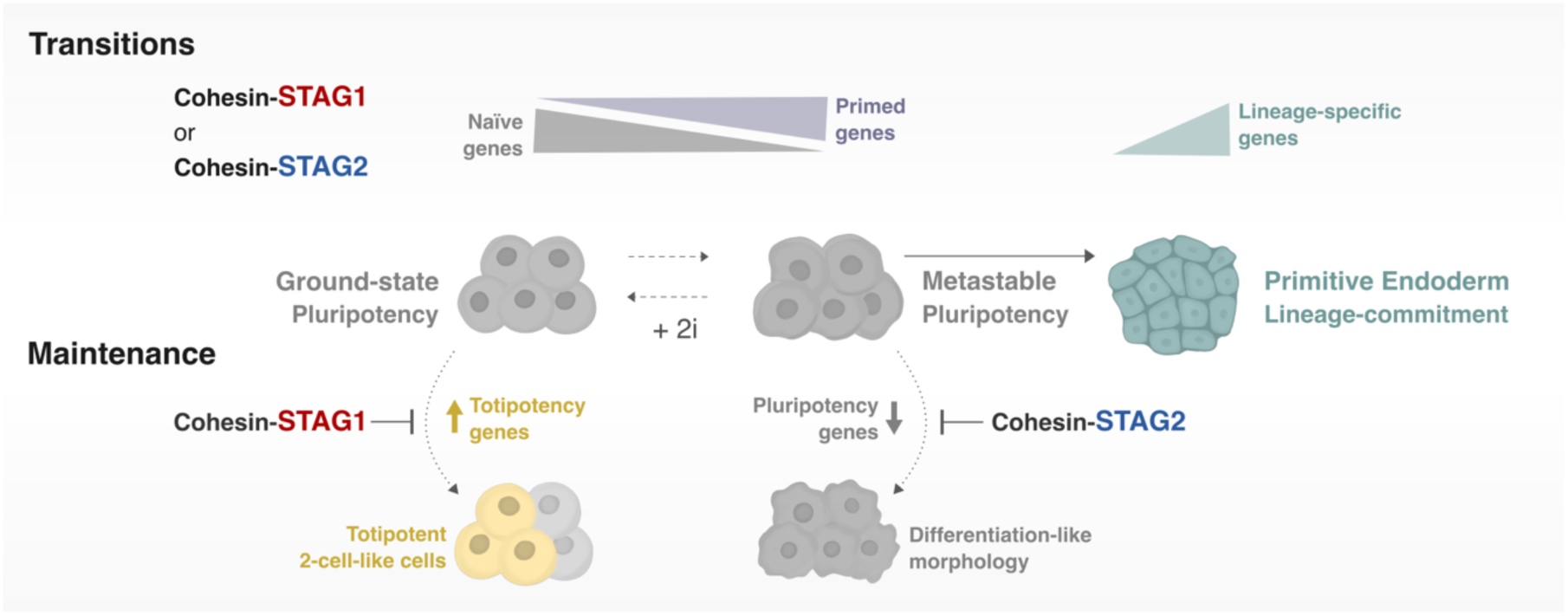
Contributions of cohesin-STAG1 and cohesin-STAG2 to pluripotency transitions and maintenance. Any of the two cohesin variants is sufficient to for mESC to transition between the ground-state and metastable states of naïve pluripotency as well as to commit to Primitive endoderm lineage. However, they have some distinct contributions to maintenance of the pluripotent state. While cohesin-STAG1 prevents return to totipotency, cohesin-STAG2 restrains exit towards commitment.

### Differentiation

We here have shown that differentiation of mESC to primitive endoderm can occur in the absence of either STAG variant, or after partial reduction of total cohesin. Cells in all tested conditions displayed the expected morphology and appropriate expression of major lineage markers, consistent with efficient acquisition of the PrE identity. While it has been proposed that cohesin may be particularly important to activate transcriptional programs during differentiation, few studies have addressed this issue outside the hematopoietic system, in part due to the essential role of cohesin for cell proliferation (Cuartero et al. 2018; Sasca et al. 2019; Minderjahn et al. 2022). A recent report tested instead acute depletion of NIPBL and found that differentiation towards neural precursor cells (NPCs) or astrocytes can proceed with only modest dysregulation of a subset of differentiation-specific genes (Hansen et al. 2026). Moreover, all differentiated cell types were present in adequate proportions in gastruloids made from NIPBL-depleted mESC. In another study, acute depletion of RAD21 in naïve mESC entering G1 in the presence of strong differentiation signals (Activin and FGF2) led to a dramatic loss of chromatin architecture but only modest effects in expression of genes involved in differentiation, with a strong enrichment for transcription factors, suggesting an auxiliary rather than determinant role of cohesin (Lee et al. 2026). Despite this evidence, the embryonic lethality of STAG1-null and STAG2-null mice together with the developmental disorders associated with mutations in cohesin-related genes, including STAG1, STAG2 and NIPBL, underscores the importance of cohesin function during development (Remeseiro et al. 2012; De Koninck et al. 2020; Kline et al. 2018; Piché et al. 2019; Solé-Ferran and Losada 2025). The relevance of cohesin is most likely specific to the nature of the transcriptional and epigenetic context of each cell type or transition. Understanding these specificities will require analyzing not only cell-intrinsic transcriptional programs, but also the three-dimensional cellular and tissue environment in which differentiation takes place.

### Pluripotency

Acute depletion of STAG2 in metastable mESC led to downregulation of *Nanog* and *Klf4,* and cell colonies displayed a differentiation-associated, flattened morphology. Strikingly, STAG2 was dispensable for pluripotent gene expression in ground-state mESC. The 2i conditions shield mESC from an acquisition of a primed state by reinforcing the core pluripotency network and inhibiting differentiation-associated signaling pathways (Marks et al. 2012; Hackett and Azim Surani 2014). In this context, the strong activation of the OCT4/ SOX2 network likely provides sufficient transcriptional stability to maintain naïve pluripotency. In contrast, mESC in the metastable state are exposed to competing signals, and cohesin-STAG2 may then be required for sustaining the high expression of pluripotency-associated genes. These findings are consistent with our previous report showing that siRNA-mediated depletion of cohesin-STAG2 in mESC growing in serum-LIF alters transcription of both superenhancer-connected genes, predominantly involved in pluripotency, and Polycomb-repressed lineage-specifying genes (Cuadrado et al. 2019). Another study characterizing *Stag1* KO and *Stag2* KO clones also reported a specific role of STAG2 in promoting transcription of genes within superenhancer domains (Arruda et al. 2020). This is unlike the *Stag2* KO clone characterized here, which was generated in a different mESC strain, and can be cultured both in ground-state and metastable conditions and maintains appropriate expression levels of *Nanog* and *Klf4*. We suspect that the requirement for cohesin-STAG2 in this clone has been bypassed after long-term loss through compensatory mechanisms.

### Totipotency

We have identified upregulation of totipotency and minor-ZGA genes after STAG1 or STAG2 depletion, although an increase in the percentage of 2CLC was only scored in the absence of STAG1. Previous reports have involved cohesin and CTCF in the control of 2CLC emergence (Zhu et al. 2021; Olbrich et al. 2021; Carey et al. 2026; Pezic et al. 2023). Reduced cohesin/CTCF binding in mESC has been suggested as a mechanism to decrease insulation and facilitate interactions between 2C enhancers and promoters that are otherwise restricted by TAD boundaries (Zhu et al. 2021). Cohesin-STAG1 is more important than cohesin-STAG2 for maintenance of TAD boundaries, which could explain why its depletion in mESC favors emergence of 2CLC (Kojic et al. 2018). The lag between protein degradation – of cohesin or its regulators – and activation of totipotency-associated genes may reflect a requirement for intermediate regulatory events linking changes in chromatin organization to the activation of the 2C-like transcriptional program. At least part of these changes may affect perinucleolar heterochromatin, facilitating the induction of DUX, a transcription factor that sustains the 2C-transcriptional program (Xie et al. 2022; De Iaco et al. 2017; Carey et al. 2026). Indeed, a role of STAG1 in supporting nucleolar integrity and repressing totipotency has been previously reported (Pezic et al. 2023). Nevertheless, the fact that upregulation of minor-ZGA genes is observed upon loss of cohesin-STAG1, cohesin-STAG2, NIPBL, WAPL or CTCF suggests that this program is particularly sensitive to changes in cohesin dynamics. Understanding additional requirements driving 2CLC emergence when different cohesin subunits and regulators are depleted and the molecular mechanisms underlying this process requires further investigation.

In conclusion, our findings highlight the differential contributions of cohesin variants to the maintenance of cellular identity and underscore the importance of cellular context in determining the consequences of cohesin perturbation.

## Materials and Methods

### Cell lines and culture conditions

R1 129 murine male ESC (129/Sv background; CNIO Mouse Genome Editing Unit) were grown in Dulbecco’s Modified Eagle’s Medium (DMEM, Sigma, D5671) supplemented with 15% inactivated fetal bovine serum (FBS, Biowest, S1810), Glutamax (Gibco, 35050-061), non-essential amino acids (NEAA, Gibco, 11140050), 50μM 2-mercaptoethanol (Gibco, 31350010), 50U/mL penicillin-50mg/mL streptomycin and LIF (produced in house) on 0.1% gelatin-coated plates. For ground-state pluripotency conditions, 1μM PD0325901 (MEK inhibitor, Axon Medchem 1408) and 3μM CHIR99021 (GSK3 inhibitor; Axon Medchem 1386) was added to media, also known as “2i”. Cells were incubated under 90% humidity at 37°C and 5% CO_2_. Cells were treated with 150nM dTAG^V^-1 (Tocris, 6914) for FKBP-tagged protein depletion. For ground-state pluripotency maintenance or ground-state to metastable transition experiments, cells were grown in 2i-containing medium for one week before starting the experiment.

### Genome editing

To obtain KO cell lines, plasmids carrying Cas9 and gRNA were transfected to mESC (Supplementary Table 1 and Supplementary Table 2). For dTAG cell lines, one plasmid carrying Cas9 and gRNA was transfected together with a plasmid that provided the repair template with the corresponding insertion. In brief, 500,000 mESC were transfected with 2μg of plasmid DNA using lipofectamine 2000 transfection reagent (Invitrogen, 11668019) in Gibco OptiMEM I reduced serum media. After 48 hours, antibiotic was aded (10 μg/mL Blasticidin or 1μg/mL Puromycin). Surviving cells were seeded at very low density for clonal selection and analyzed by immunoblotting.

### Whole cell extract preparation and immunoblots

Cells were collected, washed with PBS and resuspended in SDS-PAGE loading buffer (125mM Tris-HCl pH 6.8, 20% glycerol, 2% SDS, 0,2% bromophenol blue, supplemented with fresh 100mM DTT) at 10^4^ cells/μL. Cells were then sonicated with one pulse of 20 seconds at 40% amplitude in Branson Digital Sonifier. Alternatively, cells were resuspended with RIPA buffer for 30 minutes on ice. Upon centrifugation at 14000xg, supernatant was collected, and protein concentration was measured with DC Protein Assay (Bio-Rad). SDS-PAGE, protein transfer to nitrocellulose and immunoblots were performed using standard protocols. Primary and secondary antibodies used are listed in Supplementary Table 3.

### Analysis of total and chromatin-bound proteins by flow cytometry

Flow cytometry analysis of chromatin-bound proteins (Chromoflow, Figure 4B-C) was performed as described (Alonso-Gil and Losada 2023b). In brief, cells were collected, washed with PBS, pre-extracted for 5 minutes with a low salt extraction buffer (0.07% Igepal CA-630, 10mM NaCl, 5mM MgCl2, 0.1mM PMSF, 10mM Potassium Phosphate buffer pH 7.4) and fixed in 1% paraformaldehyde (PFA) for one hour on ice. To measure total protein levels (Figure 1D-G and Supplementary Figure 2), cells were fixed with 70% ethanol for at least two hours at -20°C. To compare distinct conditions and eliminate antibody staining variability, four different samples were barcoded with increasing concentrations of Succinimidyl Ester-Pacific Blue (Molecular Probes, P10163) for 30 minutes in the dark and pooled together prior to antibody staining. Barcoded samples were blocked in flow buffer (4% non-fat milk, 0.1% Igepal CA-630, 6.5mM Na_2_HPO_4_, 1.5mM KH_2_PO_4_, 2.7mM KCl, 137mM NaCl, 0.5 EDTA pH 7.5) for 5 minutes and incubated with primary and secondary antibodies for 1 hour or 30 minutes, respectively (Supplementary Table 3). DNA staining was performed overnight with 100nM ToPro3 in PBS. Flow cytometry analysis was performed on a BD LSRII Fortessa flow cytometer using BD FACSDiva software. Data were analyzed with FlowJo software v10.90.0 (Tree Star Inc.). For fair analysis of distinct conditions, after single-cell gating, 10,000 cells from each barcoded condition were exported, compared and plotted. For data quantification, scale values were exported for each condition, and the fold change of the median was calculated.

### Cell cycle and proliferation analyses

For cell cycle analyses, 250μM IdU (Sigma, I7125) was added to the culture medium 30 min before harvesting. Cells were collected, washed with PBS and fixed with 70% ice-cold ethanol overnight. DNA was denatured by incubation in 3N HCl for 20 minutes at room temperature. IdU was immunolabelled with FITC-BrdU Flow Kit (BD Biosciences, 559619) for 1 hour at 37°C. DNA was stained with 50μg/mL propidium iodide (Sigma, P4864) in the presence of 10μg/mL RNase A (Qiagen). Cells were processed in a FACS Canto II cytometer (BD biosciences).

CFSE CellTrace (ThermoFisher, C34554) proliferation assay was performed according to manufacturer instructions. On day 0, 3x10^6^ cells were labelled with 5μM CFSE CellTrace for 20 minutes at 37°C in suspension. Then, 250,000 cells/well were seeded in 6-well plates. Unstained and stained controls were collected at day 0, and each condition was collected at the same time the following 3 days. Harvested cells were fixed with 2% PFA for 20 minutes and stored at 4°C until flow cytometry analysis in a FACS Canto II cytometer (BD biosciences). Alternatively, cell proliferation was measured by manual counting. On day 0, around 80,000 cells were plated in 6-well plates and subsequently harvested by trypsinization after 24, 48 or 72 hours and counted using a Neubauer hemocytometer. Triplicates for each timepoint were counted, the mean was calculated and normalized to that of day 1.

### Primitive Endoderm differentiation

Primitive Endoderm differentiation was performed as previously described with minor changes (Anderson et al., 2017). Around 100,000 cells were plated in 0.1% gelatin-coated MW6-plates. The medium was replaced after 20 hours with PrE-differentiating medium consisting of RPMI 1640 (Gibco, 11340892), Glutamax (Gibco, 35050-061), 50U/mL penicillin-50mg/mL streptomycin and supplemented with B27 minus insulin (Gibco, A1895601), 4ng/mL of Activin A (PeproTech, 20-14P-10UG), 3μM CHIR99021 (GSK3 inhibitor; Axon Medchem 1386) and LIF (produced in house). Medium was changed every other day. By day 9, cells were split 1:3 into 0.1% gelatin-coated MW6 plates and maintained in culture until day 13. To assess differentiation, cells were collected at day 0 and day 13 of the protocol, washed with PBS and stained with PDFGRA antibody for 30 minutes on ice. After washing, cells were resuspended with PBS + 1μg/mL DAPI and analyzed in a FACS Canto II cytometer (BD biosciences). All flow cytometry data were analyzed with FlowJo software v10.90.0 (Tree Star Inc.).

### qPCR RNA expression analysis

Total RNA was extracted using the NZY Total RNA Isolation kit (NZYTech, MB13402) and cDNAs were prepared according to manufacturer’s instructions using the Superscript II reverse transcriptase (Invitrogen, 1864-014). qPCR was performed in triplicates using the SYBR Green PCR Master Mix (Applied Biosystems, 10187094) and an Abi Prism® 7900HT instrument (Applied Biosystems®). Primers used are listed in Supplementary Table 2. Quantifications were normalized to endogenous *Gapdh*, using the ΔΔCt method.

### Gene Set Enrichment Analysis (GSEA) after cohesin perturbation

GSEA Gene Set Enrichment Analysis (GSEA, (Subramanian et al. 2005)) was used to test enrichment of pre-implantation developmental gene sets (minor ZGA, ZGA and early/late 2-cell), which were generated as described in (Solé-Ferran et al. 2026). All analyses used 1,000 permutations, the weighted statistic and meandiv normalization. For datasets providing replicate expression matrices for perturbed and control conditions, standard GSEA was run using the Signal2Noise metric with gene-set permutation; for the remaining datasets, GSEA-Preranked was used on gene lists ranked by log2(perturbed/control). Gene sets were considered significantly enriched at FDR q ≤ 0.05.

### Immunofluorescence and high-content screening confocal imaging

For immunofluorescence analysis of ZSCAN4 and MERVL-gag positive cells, 4,500 cells/well were seeded in 0.1% gelatin-coated 96-well plates (Greiner Bio-One, 655087) and cultured for 3 days in the presence of DMSO/dTAG^v^-1. As control, wildtype mESC were treated with 50mM or 75mM NaCl for 18 hours followed by 6 hours of recovery period (as described in Canat et al., 2023) before fixation. Cells were fixed with 2% PFA for 15 minutes at RT, followed by 5 minutes of permeabilization with PBS-0.2% Triton X and blocked for 30 minutes with PBS-3% BSA. Antibodies used are listed in Supplementary Table 3. DAPI (1µg/mL) was used for nuclear staining and wheat germ agglutinin (WGA; W11261, 5µg/mL) for staining of cell membrane. Images were acquired with Opera Phenix Plus High Content Screening system (Perkin Elmer) using a 20x objective; at least 20 images were acquired per well.

Image analysis was performed using Harmony 5.1. (PerkinElmer) with a custom-built image analysis pipeline. For all ZSCAN4 analyses, DAPI staining was used for nuclear segmentation. For MERVL-gag analysis, in experiment #1, nuclei were identified based on DAPI staining, and a perinuclear ring was subsequently generated by expanding the nuclear region to quantify cytoplasmic MERVL-gag signal. For experiments #2-4, a Cellpose model was trained using DAPI (nuclear) and WGA (cell membrane) staining to enable whole-cell segmentation. MERVL-gag fluorescence intensity was then quantified within the segmented cell regions.

### Statistics

GraphPad Prism version 10.4.1 (GraphPad Software Inc) was used for all statistical analyses, with specific tests and parameters indicated in Figure legends.

## Acknowledgements

This research has been supported by grants PID2022-139333NB-I00 and PID2025-172571NB-I00 from the Agencia Estatal de Investigación (AEI/10.13039/501100011033), Ministerio de Ciencia, Innovación y Universidades (MICIU) and cofounded by the European Regional Development Fund (ERDF-EU). M.S.F. has been awarded predoctoral contracts from Comunidad de Madrid (PEJD-2019-PRE/BMD-15214) and AEI (PRE2020-091957). S.A. has been supported by the Ramón y Cajal program of MICIU, and the European Social Fund (RYC-2018-025002-I). The di Croce laboratory acknowledges also support of MICIU through the Centro de Excelencia Severo Ochoa program (CEX2020-001049-S, MCIN/AEI /10.13039/501100011033), and of the Generalitat de Catalunya through the CERCA program.

## Author contributions

MS-F designed, performed and analyzed all experiments. DG-Ll performed RNA-seq data analyses. M.R. provided technical assistance with western blot analyses. AC initiated the project and provided advice throughout its execution. Initial differentiation experiments to PrE were performed in LdC lab under the supervision of SA. AL obtained funding, supervised the study and wrote the manuscript with MS-F and contributions from all other authors.

## Competing interests

The authors declare no competing interests.

---

**Supplementary Figure 1.**
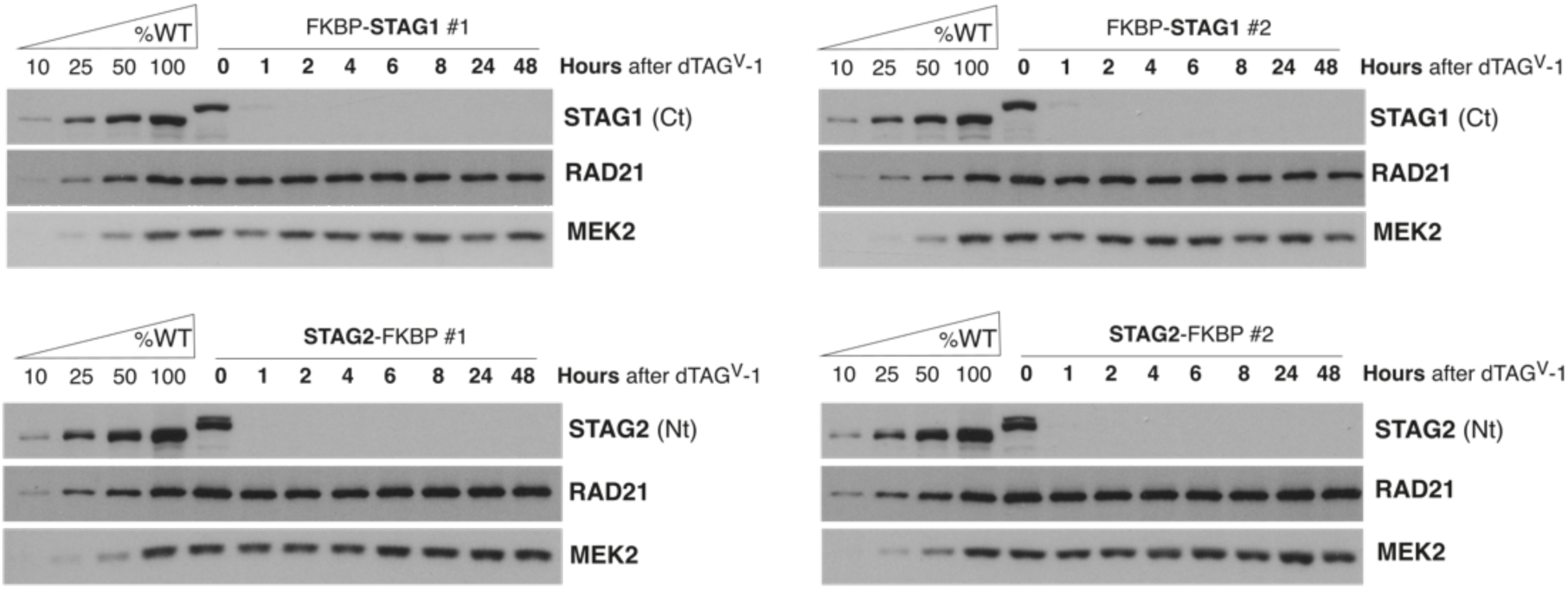
Fast and extensive degradation of STAG1 and STAG2 using degrons. Immunoblots of whole cell extracts of two clones each of FKBP-STAG1 and STAG2-FKBP cells to monitor depletion of the corresponding protein at different times after addition of 150nM dTAG^v^1. Decreasing amounts of an extract prepared from wildtype mESC (WT) were analyzed in the same gels to allow quantitative comparisons. STAG1 C-terminal (Ct) and STAG2 N-terminal (Nt) antibodies were used to avoid interference with the tag. MEK2 was used as a loading control.

**Supplementary Figure 2.**
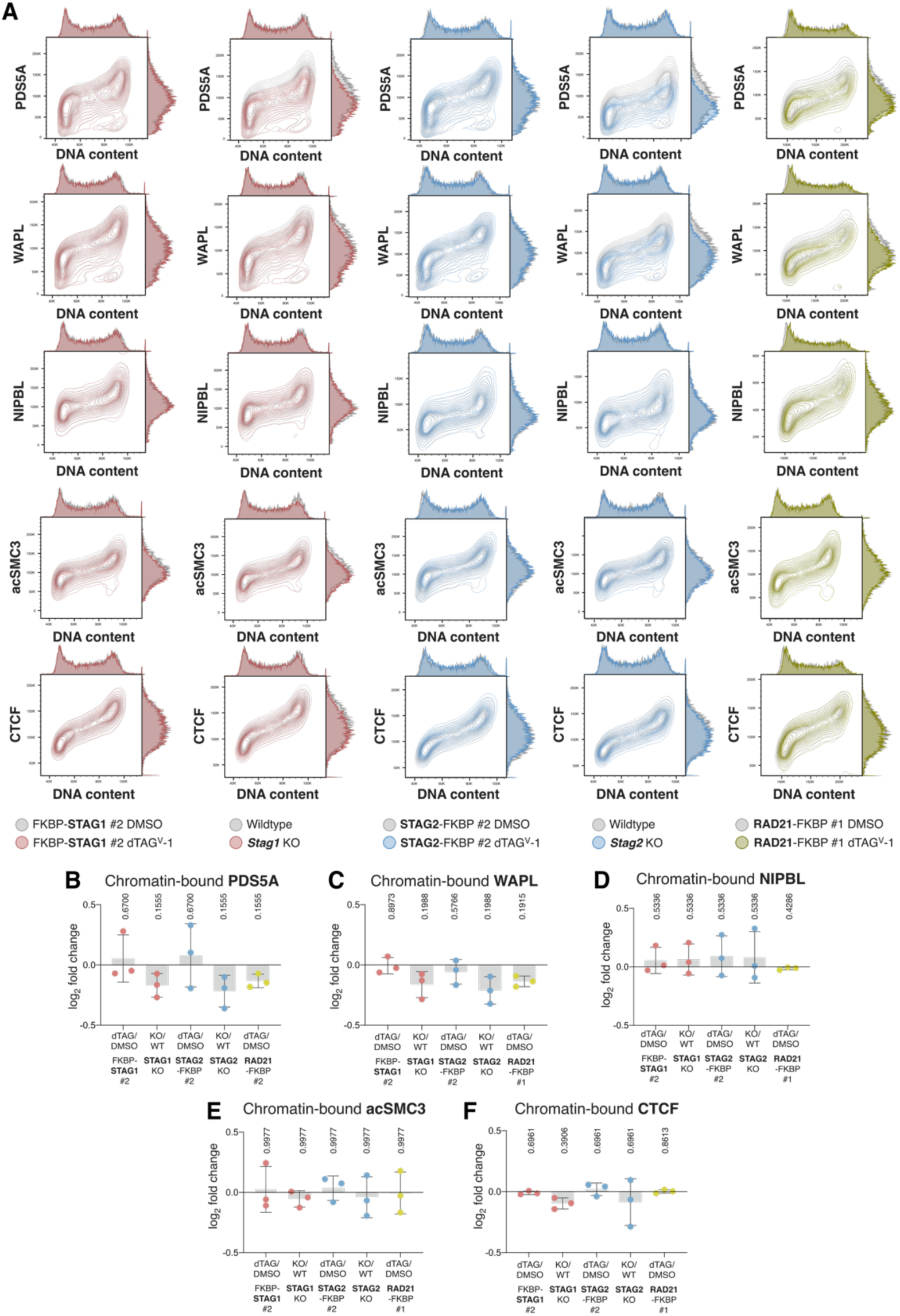
No changes in chromatin-bound levels of Cohesin regulators upon STAG1 or STAG2 depletion. **A.** Contour plots of chromatin-bound levels of the indicated cohesin regulators along the cell cycle in the different clones, assessed by Chromoflow. For FKBP clones, DMSO-treated cells (in grey) are compared to dTAG^v^-1-treated cells (colored). For KO clones (colored) the comparison is to wildtype mESC (in grey). B-F. Quantification of at least three independent experiments as in (A) for PDS5A (B), WAPL (C), NIPBL (D), acSMC3 (E) and CTCF (F). The log2 fold change relative to controls (dTAG^v^-1- vs DMSO-treated or KO vs WT) was calculated from the median fluorescence intensity for each independent experiment (dot), the bar represents the mean and the whiskers the standard deviation. To assess statistical significance, multiple paired-tests were performed on the raw median fluorescence intensity values, with correction for multiple comparisons using the two-stage Bejamini, Krieger and Yekutieli false discovery rate procedure.

**Supplementary Figure 3.**
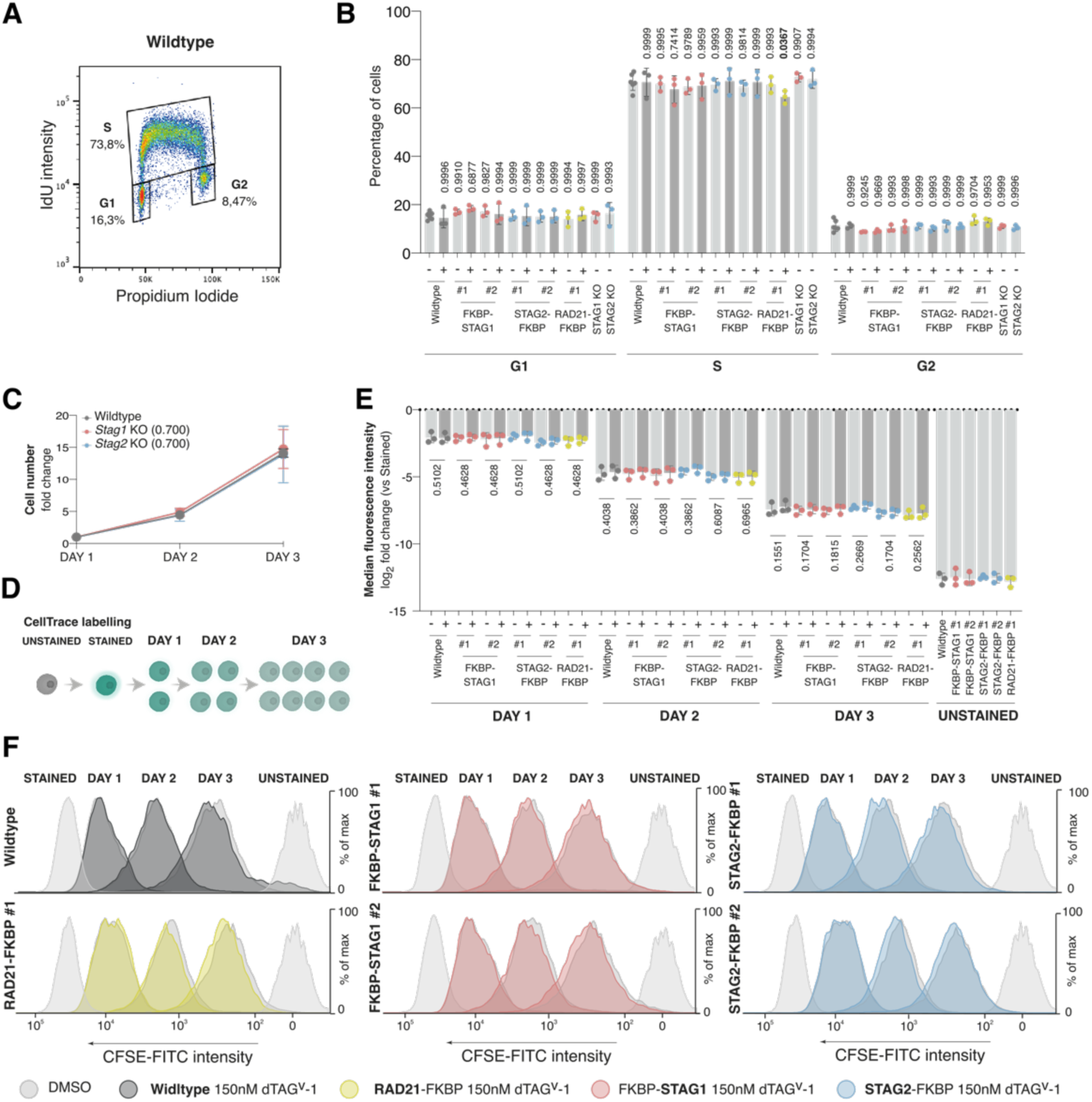
No alterations in cell cycle profiles or proliferation rates after STAG1 or STAG2 depletion. A. Representative cell cycle profile of wildtype mESC, exposed to a 30-min pulse of BrdU before harvesting. Gated G1, S and G2 populations are indicated. B. Percentage of G1, S and G2 cell populations from the distinct clones, gated as in (A). Each dot corresponds to one experiment; the bar represents the mean and whiskers the standard deviation. P-values were calculated using a two-way ANOVA followed by Dunnett’s correction for multiple comparisons. For every cell cycle phase, each condition was compared to wildtype untreated control. C. Growth curves of *Stag1* and *Stag2* KO clones compared to wildtype mESC. Fold change was calculated relative to cell number at DAY 1. Three independent experiments with triplicates were performed for each clone. The dots represent the mean of all three experiments and whiskers the standard deviation. P-values for DAY 3 were calculated using an unpaired t-test D. Illustration of the experimental design to measure cell proliferation by flow cytometry. Cells stained with CellTrace-CFSE at DAY 0 are followed for three consecutive days as fluorescence intensity is halved with each division. E. Quantification of the three CellTrace experiments performed (representative images in F). The log2 fold change was calculated from the median fluorescence intensity relative to DAY 0 stained controls for each clone. Each dot corresponds to one experiment; the bar represents the mean and the whiskers the standard deviation. To assess statistical significance, multiple paired-tests were performed on the raw median fluorescence intensity values, with correction for multiple comparisons using the two-stage Bejamini, Krieger and Yekutieli false discovery rate procedure. Comparisons were made between DMSO-and dTAG^v^-1 treated clones. F. Representative histogram plots, normalized to their higher peak (mode), of CFSE-FITC intensity in the indicated clones over three days, comparing DMSO- and dTAG^v^-1-treated cells. Unstained and Stained controls were collected and fixed on the day of seeding for each of the clones analyzed.

**Supplementary Figure 4.**
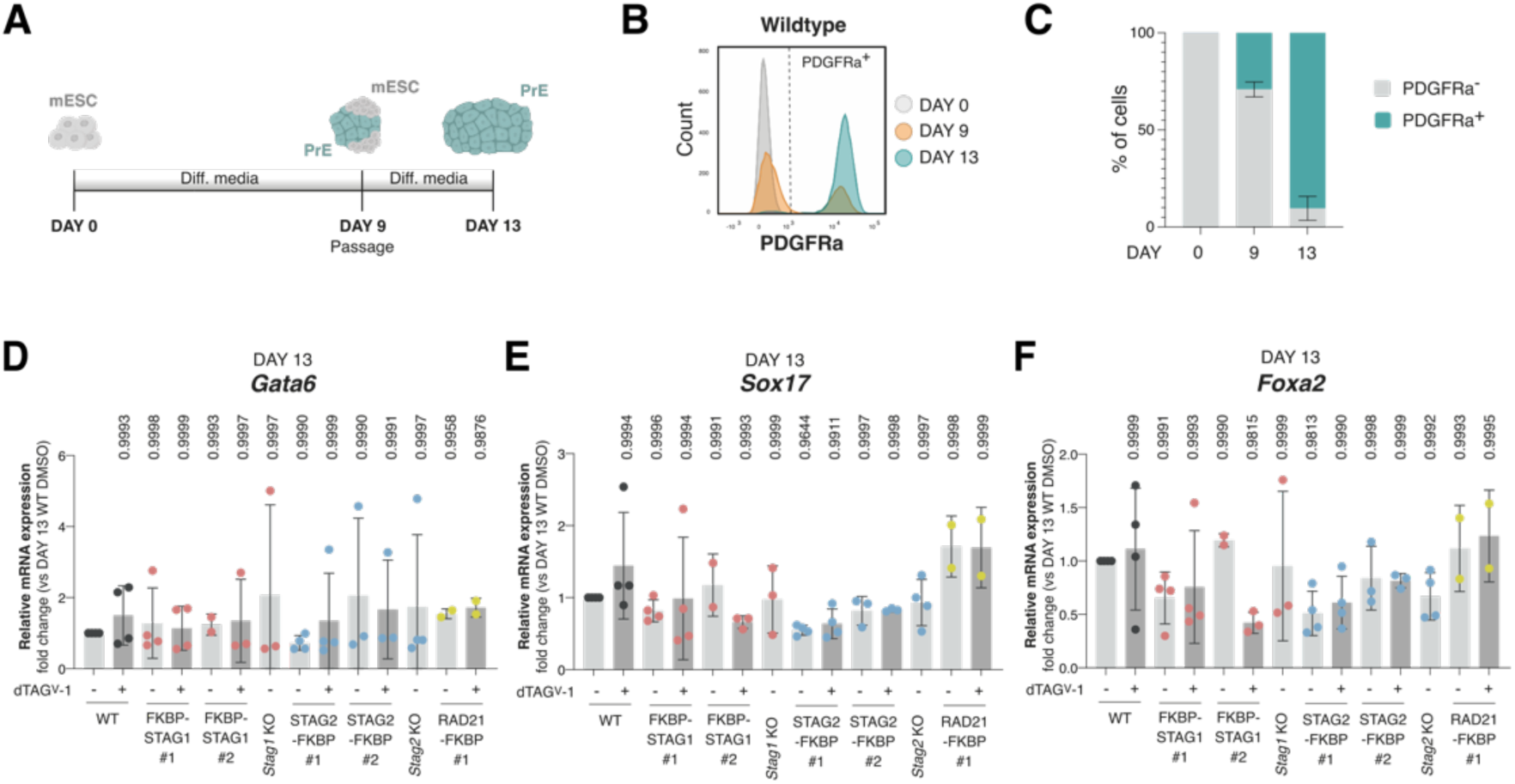
Proper acquisition of Primitive Endoderm fate in the absence of STAG1 or STAG2. A. Schematic illustration of the experimental design. mESC are seeded and cultured in differentiation medium for 9 days, when they are passaged and cultured for 4 more days until a pure primitive endoderm population is obtained. B. PFGFRa intensity measured at the indicated days of differentiation for wildtype mESC. C. Quantification of the percentage of PDGFRa positive cells from (B) in three independent experiments. Whiskers indicate standard deviation. D-F. Fold changes in *Gata6* (D), *Sox17* (E) and *Foxa2* (F) transcripts in DMSO- or dTAG-treated clones at DAY 13 of differentiation. Each sample was first normalized by *Gapdh* expression and then to expression levels in DMSO-treated wildtype differentiated cells. Each dot corresponds to an independent experiment, the bar represents the mean and the whiskers the standard deviation. P-values were calculated using a two-way ANOVA followed by Dunnett’s multiple-comparisons test on the raw ΔCt values. Each condition was compared to DMSO-treated wildtype differentiated cells.

**Supplementary Figure 5.**
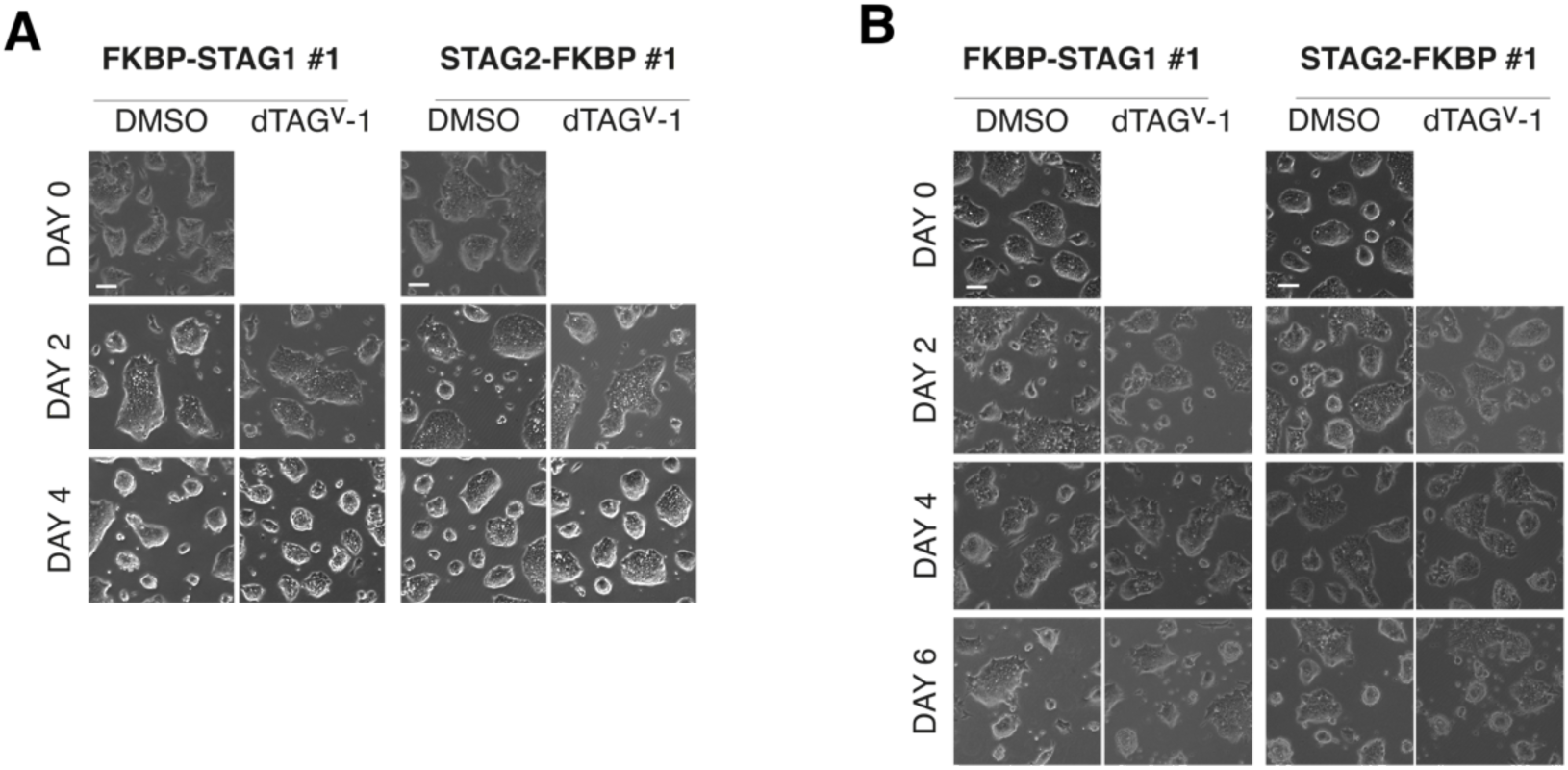
Successful transitions between ground and metastable pluripotency states in the absence of STAG1 or STAG2. A-B. Representative images of cells of the indicated clones during the transitions between metastable pluripotent and ground sates upon addition (A) or retrieval (B) of 2i, as in Figure 3B and Figure 3G, respectively. Scale bar, 20µm.

**Supplementary Figure 6.**
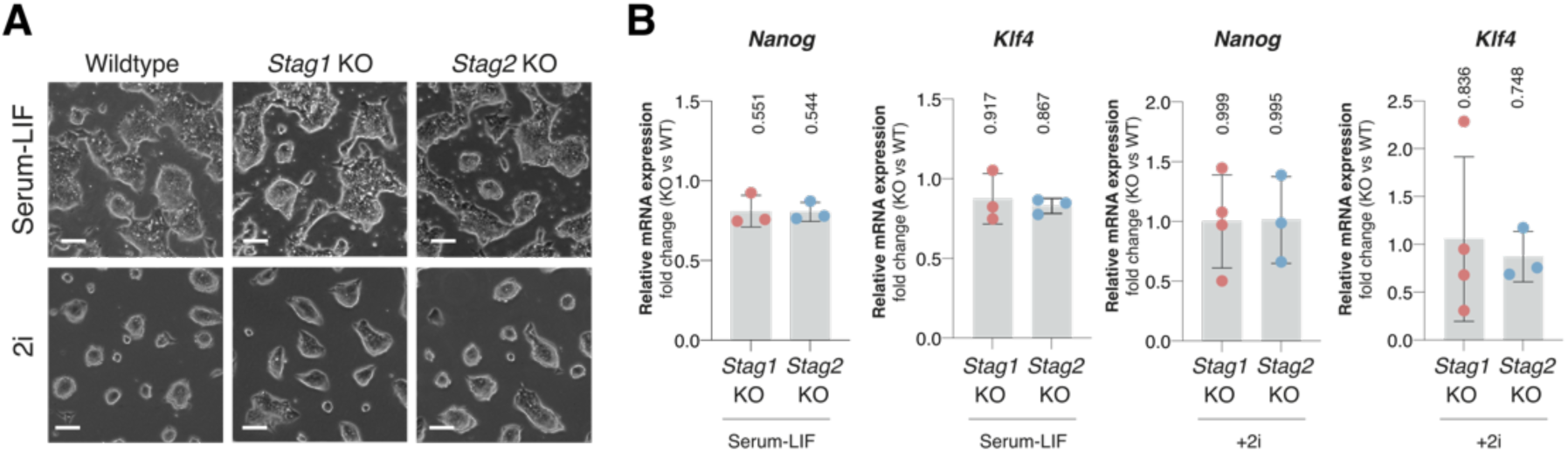
Proper maintenance of metastable and ground-state pluripotency in *Stag1* and *Stag2* KO cells. A. Representative images of the indicated clones cultured in serum/LIF (metastable) or in the presence of 2i (naïve). Scale bar, 20µm. B. Fold changes in *Nanog* and *Klf4* transcripts in *Stag1* and *Stag2* KO clones cultured as in (A). Each sample was first normalized by *Gapdh* expression and then to expression levels in wildtype mESC. Each dot corresponds to one experiment; the bar represents the mean and the whiskers the standard deviation. P-values were calculated using a regular one-way ANOVA followed by Dunnett’s multiple-comparisons test on the raw ΔCt values. Each condition was compared to wildtype mESC.

**Supplementary Table 1.** Plasmids used in this study.

| Plasmid | Use |
|---|---|
| pX459-MSF9 | To obtain *Stag1* KO cells. gRNA + Cas9 expressing vector. Puromycin resistance. |
| pX459-MSF12 | To obtain *Stag2* KO cells. gRNA + Cas9 expressing vector. Puromycin resistance. |
| pX459-MSF5 | To target *Stag1* N-ter. gRNA + Cas9 expressing vector. Puromycin resistance. |
| pX459-MSF9 | To target *Stag2* C-ter. gRNA + Cas9 expressing vector. Puromycin resistance. |
| pEN243-1082 | To target *Rad21* C-ter. gRNA + Cas9 expressing vector. Puromycin resistance. |
| pMSF1 | For STAG1 N-terminal tagging with FKBP. |
| pMSF2 | For STAG2 C-terminal tagging with FKBP. |
| pMSF13 | For RAD21 C-terminal tagging with FKBP. Derived from pEN862. |

**Supplementary Table 2.**
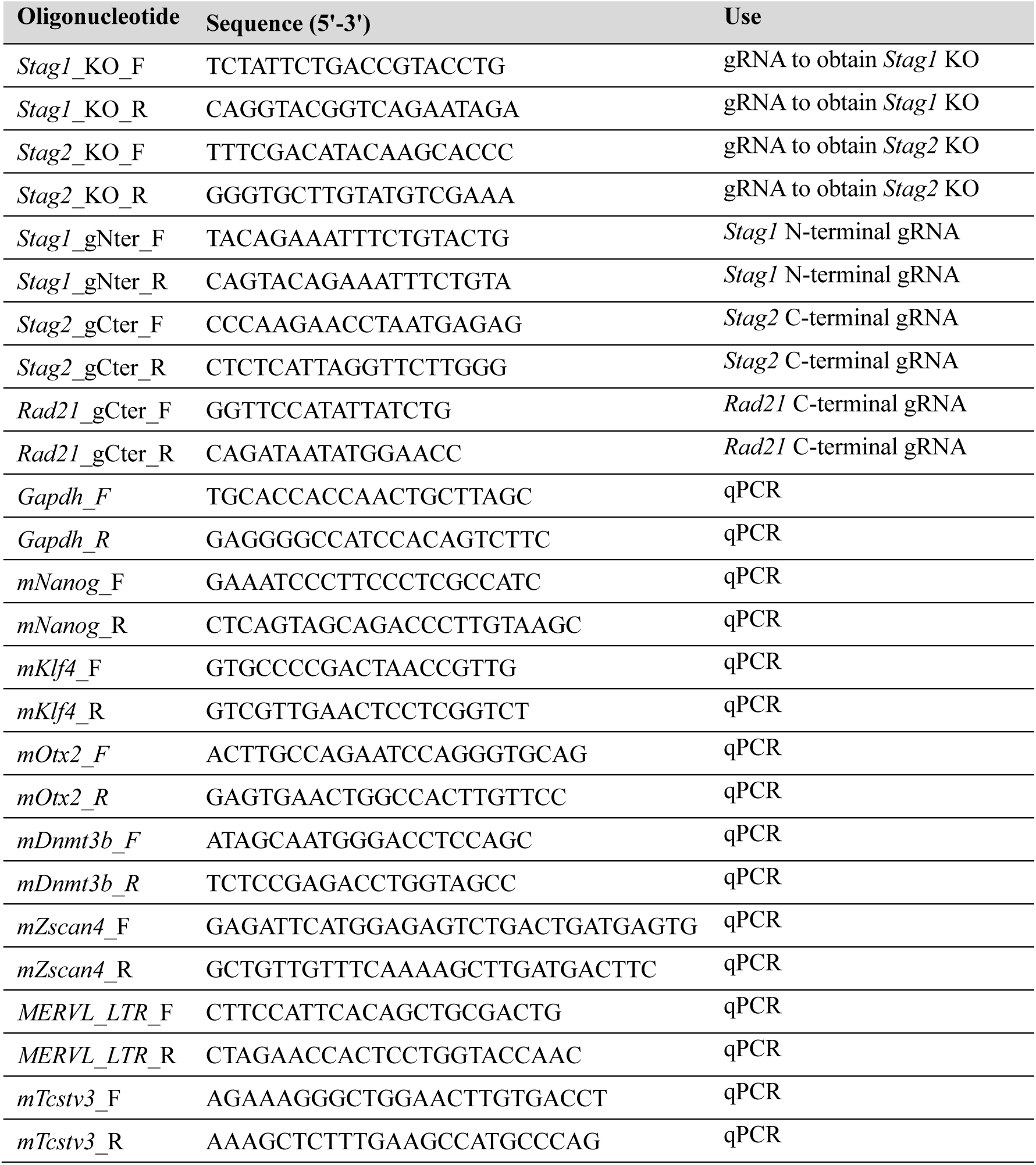
Oligonucleotides used in this study.

**Supplementary Table 3.** Antibodies used in this study.

| **Antibody** | **Reference** | **Use\*** |
|---|---|---|
| MEK2 mouse monoclonal | M24520 (BD Biosciences) | 1:1000 for IB |
| TUBULIN rat monoclonal | YL1/2 | 1:40000 for IB |
| RAD21 rabbit polyclonal | Carretero et al., 2013 | 1µg/mL for IB |
| STAG1 rabbit polyclonal (Ct) | custom made, Remeseiro et al., 2012 | 1µg/mL for IB, 4µg/mL for FC |
| STAG1 rat monoclonal (Nt) | Custom made, Kojic et al., 2018 | 1:10 diluted supernatant for IB |
| STAG2 rabbit polyclonal (Nt) | Custom made, Giménez-Llorente et al., 2024 | 1µg/mL for IB, 4µg/mL for FC |
| RAD21 mouse monoclonal | Merck, 05-908 | 2µg/mL for FC, 2µg/mL for IB |
| Acetyl SMC3 mouse monoclonal | Custom made, Nishiyama et al., 2010 | 1:50 for FC |
| WAPL rabbit polyclonal | Custom made, Kojic et al., 2018 | 4µg/mL for FC |
| NIPBL mouse monoclonal | Santa Cruz Biotechnology, sc-374625 | 2µg/mL for FC |
| PDS5A rabbit polyclonal | Custom made, Carretero et al., 2013 | 2µg/mL for FC |
| CTCF mouse monoclonal | Santa Cruz Biotechnology, sc-271474 | 2µg/mL for FC |
| NANOG rabbit polyclonal | Abcam, AB5731 | 1:1000 for IB, 1:200 for FC |
| NANOG rat monoclonal | eBioscience, 14-5761 | 1:200 for FC |
| PDGFRA APC conjugated mouse monoclonal | eBioscience, 12-1401-81 | 1:300 for FC |
| ZSCAN4 rabbit polyclonal | Sigma, AB4340 | 1:1500 for IB, 1:1000 for IF |
| MERVL-gag rabbit polyclonal | Novus Biologicals, NBP2-66963 | 1:1000 for IF |
| Cy3-conjugated donkey anti-rabbit | Jackson Immunoresearch, 711-095-152 | 1:200 for IF, 1:400 for FC |
| Alexa Fluor 488-conjugated goat anti-mouse | Abcam, AB_2340846 | 1:400 for FC |

\*FC, flow cytometry; IB, immunoblot; IF, immunofluorescence

## Notes

### Competing Interest Statement

The authors have declared no competing interest.

